# Can processing of face trustworthiness bypass early visual cortex? A transcranial magnetic stimulation masking study

**DOI:** 10.1101/649350

**Authors:** Shanice E. W. Janssens, Alexander T. Sack, Sarah Jessen, Tom A. de Graaf

**Affiliations:** Section Brain Stimulation and Cognition, Department of Cognitive Neuroscience, Faculty of Psychology and Neuroscience, Maastricht University, Maastricht, the Netherlands; Maastricht Brain Imaging Centre (MBIC), Maastricht University, Maastricht, the Netherlands; Department of Neurology, University of Lübeck, Lübeck, Germany

**Author notes:** Corresponding author: Contact details, Oxfordlaan 55, 6229 EV, Maastricht, the Netherlands, Section Brain Stimulation and Cognition, Department of Cognitive Neuroscience, Faculty of Psychology and Neuroscience, Maastricht University, Maastricht, the Netherlands.

**Keywords:** trustworthiness, early visual cortex (EVC), transcranial magnetic stimulation (TMS) masking, unconscious processing

## Abstract

As a highly social species, we constantly evaluate human faces to decide whether we can trust someone. Previous studies suggest that face trustworthiness can be processed unconsciously, but the underlying neural pathways remain unclear. Specifically, the question remains whether processing of face trustworthiness relies on early visual cortex (EVC), required for conscious perception. If processing of trustworthiness can bypass EVC, then disrupting EVC should impair conscious trustworthiness perception while leaving forced-choice trustworthiness judgment intact. We applied double-pulse transcranial magnetic stimulation (TMS) to right EVC, at different stimulus onset asynchronies (SOAs) from presentation of a face in either the left or right hemifield. Faces were slightly rotated clockwise or counterclockwise, and were either trustworthy or untrustworthy. On each trial, participants discriminated 1) trustworthiness, 2) stimulus rotation, and 3) subjective visibility of trustworthiness. At early SOAs and specifically in the left hemifield, orientation processing (captured by the rotation task) was impaired by TMS. Crucially, though TMS also impaired subjective visibility of trustworthiness, no effects on trustworthiness discrimination were obtained. Conscious perception of face trustworthiness (captured by visibility ratings) relies on intact EVC, while forced-choice trustworthiness judgments may not. These results are consistent with the hypothesis that trustworthiness processing can bypass EVC. For basic visual features, extrastriate pathways are well-established; but face trustworthiness depends on a complex configuration of features. Its processing without EVC and outside of awareness is therefore of particular interest, further highlighting its ecological relevance.

## Introduction

When meeting someone for the first time, we immediately form an impression about whether that person is friend or foe. Solely based on facial features, a person is judged to be more or less trustworthy (Todorov, Olivola, Dotsch, & Mende-Siedlecki, 2015; Todorov, Said, Engell, & Oosterhof, 2008). An exposure of less than 100 ms is sufficient to make a trustworthiness judgment that is highly replicable and consistent across observers (Todorov, Pakrashi, & Oosterhof, 2009; Willis & Todorov, 2006). This suggests that trustworthiness judgments rely on rapid and automatic processing streams, but does not establish this processing to be unconscious. On the other hand, for a range of basic visual features, as well as more complex inputs that are ecologically relevant, behaviorally relevant processing *can* occur outside of awareness (De Gelder, Haan, & Heywood, 2001).

Although there are different theories on the neural basis of visual awareness, they seem to agree that awareness involves recurrent activity within and between stages of the visual hierarchy (Dehaene & Naccache, 2001; Lamme & Roelfsema, 2000; Tononi, 2008). As a result, a conscious percept takes time to establish. For inputs that seem particularly relevant, additional processing time should not delay our initial behavioral or emotional responses. Such relevant information can guide our behavior before, or even fully without, the onset of awareness. Given the speed and ecological relevance of trustworthiness processing, can it, too, occur outside of awareness?

Recent studies addressed this question using continuous flash suppression (CFS). In CFS, a visual stimulus is presented to one eye and rendered unconscious by simultaneously presenting dynamic stimulation to the other eye (Tsuchiya & Koch, 2004). (Un)trustworthy faces took longer to “break through” CFS (i.e., become visible) compared to neutral faces (Getov, Kanai, Bahrami, & Rees, 2015; Stewart et al., 2012). Further evidence for unconscious processing of face trustworthiness comes from subliminal priming: subliminally presented (un)trustworthy faces biased the judgment of subsequent supraliminal neutral faces (Todorov et al., 2009). On a neural level, the amygdala shows response selectivity to both conscious (Santos, Almeida, Oliveiros, & Castelo-Branco, 2016) and unconscious (Freeman, Stolier, Ingbretsen, & Hehman, 2014) face trustworthiness. In sum, both behavioral and neural studies suggest that face trustworthiness can be processed unconsciously.

We hypothesize that unconscious trustworthiness processing relies on a pathway bypassing early visual cortex (EVC), analogous to unconscious emotion processing (Gainotti, 2012). One way to address this hypothesis is to use transcranial magnetic stimulation (TMS) to disrupt EVC and thereby impair performance on visual tasks (Kammer, 2007a). That approach has successfully been used to investigate many visual functions, including letter identification (Amassian et al., 1989; Masur, Papke, & Oberwittler, 1993; Potts et al., 1998), number identification (Miller, Fendrich, Eliassen, Demirel, & Gazzaniga, 1996) and orientation discrimination (Beckers & Hömberg, 1991; Boyer, Harrison, & Ro, 2005; Kammer, Puls, Strasburger, Hill, & Wichmann, 2005). In such TMS masking studies, TMS is delivered at several stimulus-onset asynchronies (SOAs). Visual task performance is most consistently impaired when TMS is delivered to EVC at around 70–130 ms SOA (De Graaf, Koivisto, Jacobs, & Sack, 2014; Kammer, 2007a, 2007b).

Previous TMS masking studies have implemented both objective and subjective tasks to dissociate the relevance of EVC for unconscious versus conscious perception (Boyer et al., 2005; Koenig & Ro, 2018; Christensen, Kristiansen, Rowe, & Nielsen, 2008). Subjective measures such as visibility ratings capture conscious visual perception, while objective measures such as forced-choice tasks can also capture unconscious visual processing. For instance, Jolij and Lamme (2005) showed that discrimination of emotional expressions remains intact after occipital TMS, whereas performance was impaired in an objective location discrimination task on the same stimuli. In other words, unconscious emotion processing was possible without EVC. Since trustworthiness judgments are highly ecologically relevant as well, we hypothesize that processing of face trustworthiness can also occur without the involvement of EVC.

To test this, we used double-pulse TMS at different SOAs to disrupt processing in EVC. Participants performed a two-alternative forced-choice (2AFC) rotation discrimination (control) task, included to capture orientation processing and validate the neural efficacy of our TMS protocol (De Graaf & Sack, 2010, 2018). We furthermore implemented a subjective trustworthiness visibility rating task, to capture conscious trustworthiness perception, and a 2AFC trustworthiness discrimination task, to capture potentially unconscious trustworthiness processing, to directly dissociate TMS effects on the two tasks. According to our hypothesis, TMS to EVC should reduce subjective visibility of trustworthiness; but if trustworthiness processing can bypass EVC, then trustworthiness discrimination should remain intact.

## Methods

### Participants

Twenty volunteers participated in this experiment (9 males, ages 18–29). Participants were screened for transcranial magnetic stimulation (TMS) contraindications, provided written informed consent and had (corrected-to-)normal vision. The experiment was approved by the local ethical committee. Participants were compensated with either participation credits or 10 euros in vouchers per hour.

### Stimuli, Tasks, and Design

In one 2-hour session, participants performed several tasks in response to face stimuli (see Figure 1). We selected 25 male Caucasian faces from an existing database (Oosterhof & Todorov, 2008; Todorov, Baron, & Oosterhof, 2018). For each face, we used a version classified as untrustworthy (−3 SD from the average neutral face) and a version classified as trustworthy (+3 SD from the average neutral face) (Oosterhof & Todorov, 2008), leading to 50 different faces in total. We disrupted early visual cortex (EVC) of the right hemisphere with 20-Hertz double-pulse TMS at different stimulus-onset-asynchronies (SOAs). Trials without TMS served as a control condition (factor “TMS”: 50 ms, 100 ms, 150 ms, no TMS). Faces were presented in the left or right hemifield (factor “Hemifield”), varying in trustworthiness and rotation (from the vertical meridian). To prevent floor and ceiling performance, face opacity and rotation angle were individually calibrated prior to the main task. Participants performed three tasks, capturing three visual processing mechanisms:

- Trustworthiness discrimination;
- Rotation discrimination;
- Subjective rating of trustworthiness visibility.

**Figure 1.**
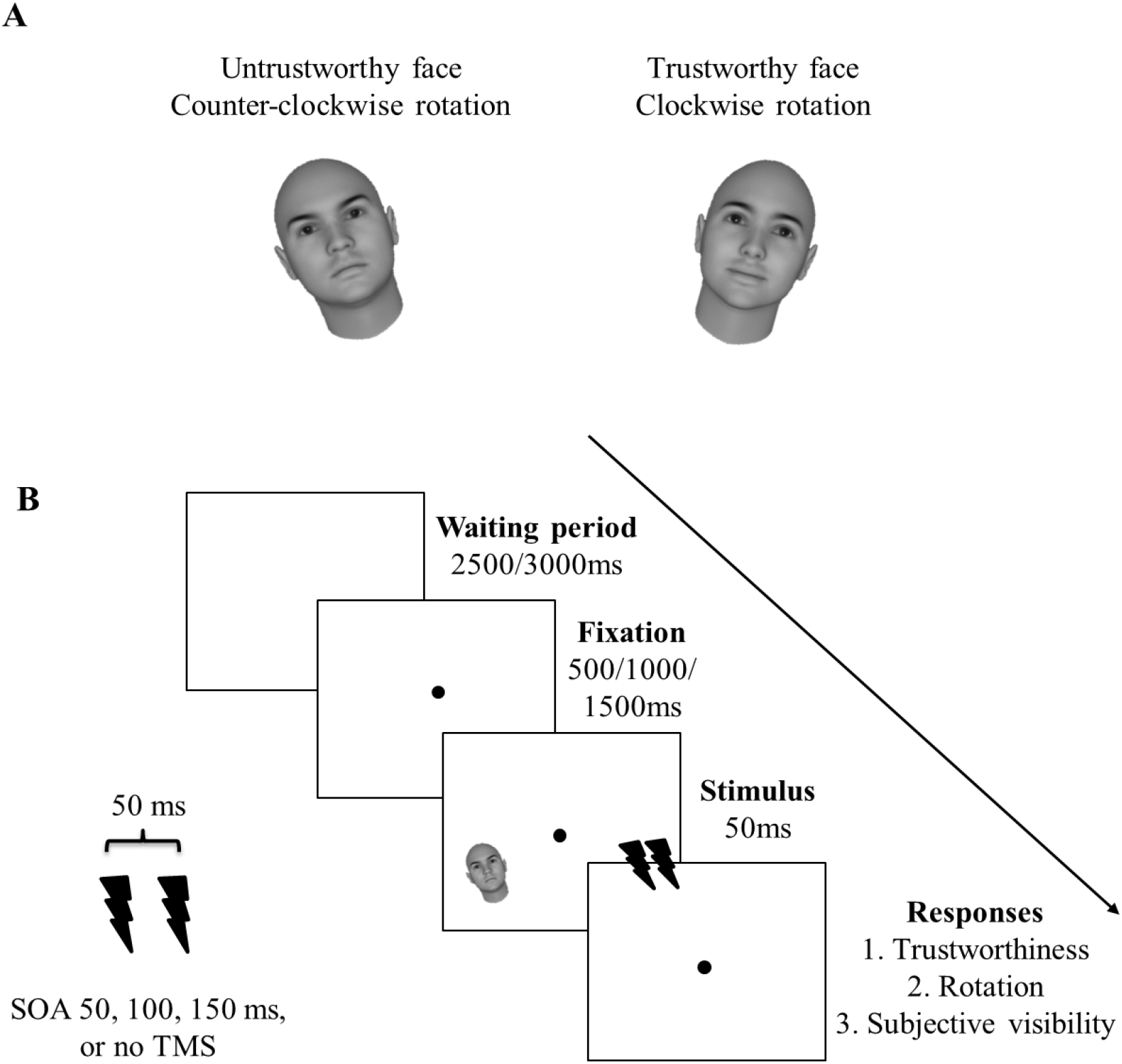
Stimuli and design. A) Example stimuli. We presented faces which are either untrustworthy or trustworthy from a stimulus set created by Oosterhof and Todorov (2008). The faces were rotated either clockwise or counter-clockwise. B) Experimental design. In each trial, either no TMS was applied, or double-pulse TMS with an inter-pulse interval of 50 ms was applied at a stimulus-onset asynchrony (SOA) of 50 ms, 100 ms, or 150 ms (factor TMS). Faces were presented either to the lower left or right of fixation (factor Hemifield). After each stimulus, participants first indicated the trustworthiness of the stimulus (2AFC: trustworthy vs untrustworthy), then the rotation of the stimulus (2AFC: clockwise vs counter-clockwise), and then the subjective visibility of the trustworthiness of the stimulus (4-point scale).

Rotation discrimination involved a two-alternative forced choice (2AFC) task, indicating by button press whether faces were rotated clockwise or counterclockwise from vertical. Performance should be impaired by TMS pulses around 100 ms, since it relies on orientation processing (De Graaf, Duecker, Fernholz, & Sack, 2015; De Graaf, Cornelsen, Jacobs, & Sack, 2011; De Graaf, Herring, & Sack, 2011; De Graaf et al., 2014; Jacobs, De Graaf, & Sack, 2014) (‘neural efficacy check’, (De Graaf & Sack, 2010, 2018)). Trustworthiness discrimination was a 2AFC task (trustworthy vs untrustworthy). Subjective trustworthiness visibility ratings were given on a 4-point scale (1 (“not perceived”) to 4 (“clearly perceived”), with 2 and 3 parametrically in between), inspired by other 4-point subjective scales (e.g., perceptual awareness scale (PAS) (Ramsøy & Overgaard, 2004)) and same as in our previous studies (De Graaf, Goebel, & Sack, 2012; De Graaf et al., 2011a; De Graaf et al., 2011b; Jacobs et al., 2014; Jacobs, Goebel, & Sack, 2012). Participants were explicitly instructed that subjective visibility ratings should reflect how well the *trustworthiness* was perceived, not the stimulus itself.

### TMS parameters

Biphasic TMS pulses were delivered using a MagPro R30 stimulator (MagVenture, Farum, Denmark) and a figure-of-eight coil (MC-B70). Stimulation intensity was 60% maximum stimulator output (MSO), unless this exceeded 200% of phosphene threshold (PT), in which case stimulation was at 200% PT (45% MSO, one case), or participants indicated discomfort in which case stimulation intensity was set to a tolerable level (five cases: 45% MSO (1), 50% MSO (2), 55% MSO (2)). PT was defined as the stimulation intensity at which phosphenes were reported in half of trials, in an informal procedure (average: 39%, range: 23–47% MSO). We failed to record phosphene perception in four participants, we recorded that eight participants saw phosphenes, and eight participants did not see phosphenes. If phosphenes could not be elicited, the coil was positioned 2 cm above and to the right of the inion. If the participant perceived phosphenes overlapping with the left stimulus location, the coil was positioned on the phosphene hotspot; the location over right EVC with lowest PT. During the main task, coil position was fixed with lateral handle orientation using a mechanical arm. On each trial, double-pulse TMS with an inter-pulse interval of 50 ms was delivered at 50 ms, 100 ms, 150 ms, or not at all (no TMS) (see Figure 1B). Note that SOA conditions connected: the second pulse for one SOA was at the same time relative to visual stimulus onset as the first pulse of the next SOA condition. The waiting period between trials was either 2.5 or 3 sec. The total number of pulses (excluding PT determination) was 360.

### Procedures

Stimuli were presented using MATLAB (The MathWorks, Inc., Natick, Massachusetts, United States) with Psychophysics Toolbox (Brainard, 1997) on a 24 inch monitor with a refresh rate of 60 Hz. Faces were 4 degrees visual angle (DVA) high and were presented in the left or right hemifield, 3 DVA diagonally below central fixation, for 50 ms. A black fixation dot of 0.3 DVA was presented in the middle of a grey screen with a background luminance of 156 cd/m2. Participants rested their heads in a chin rest, eyes 57 cm from screen, fixating continuously.

We calibrated task difficulty for both the rotation and trustworthiness task, using psychophysical staircases. We determined required face rotation (in degrees) for 80% discrimination accuracy. Quest (Watson & Pelli, 1983) was used with the following parameters: tGuess = ^10^log(4), tSD = ^10^log(10), beta = 3.5, delta = 0.01, gamma = 0.5. In each trial, after 0.5, 1 or 1.5 sec fixation, a face appeared lower left of fixation. Participants reported as quickly and as accurately as possible the (counter-)clockwise rotation of the face. The task lasted ~3 minutes and included 5 practice trials with feedback, and 50 main trials. We plotted rotation test values over trials: if staircases did not converge to a stable value they were repeated.

Another staircase determined required face opacity for 80% correct (un-)trustworthy discrimination. Staircase procedures and parameters were the same as above, but preceded by 10 practice trials in which stimuli were shown at full contrast for 500 ms. Quest here was provided with tGuess (^10^log(125)) and tSD (^10^log(255)). In four participants, the task proved too difficult, and stimulus duration was increased to 67 ms throughout the experiment.

The ‘performance check’ included 5 practice and 40 main trials, using rotation and opacity values resulting from the staircases. As in the main experiment, subjects provided three responses per stimulus: first they indicated the trustworthiness, then rotation of the face. Lastly, they rated subjective visibility of (how clearly they could perceive) the *trustworthiness* of the face on a four-point scale. As in the main experiment, response options and corresponding keys for rotation discrimination and subjective visibility ratings were prompted on screen on each trial. The performance check took 5 minutes, including one break. With now three required responses per trial, participants sometimes needed more practice. Therefore, if performance for either of the 2AFC tasks was close to floor (<55%) or ceiling (>95%), the performance check was repeated.

The main experiment was described above (see Figure 1). With factors TMS (50 ms, 100 ms, 150 ms, no TMS) and target hemifield (left, right), and 30 trials per condition cell, the task included 240 trials in total, preceded by 5 practice trials, offered 9 breaks at regular intervals, and lasted ~45 minutes.

### Analysis

Dependent variables for the 2AFC tasks were proportion correct, and average rating for the subjective trustworthiness visibility task. If participants performed too close to ceiling (>95%) or floor (<55%) in the no TMS control condition, they were excluded from the analyses *for that task*: three participants were excluded from the trustworthiness task and three from the rotation task (of those, one participant was excluded from both tasks).

The condition means of the three dependent variables were compared with two-way repeated measures ANOVAs with the factors “Hemifield” (left, right) and “TMS” (50ms, 100ms, 150ms, no TMS), in SPSS Version 24.0 (IBM Corp., Armonk, New York, United States). For completeness and comparison, we report both Greenhouse-Geisser-corrected univariate results and multivariate results. Significant interactions were followed by simple effects analyses. Additionally, TMS SOA conditions were contrasted to no-TMS in Bonferroni-corrected planned comparisons. To provide more direct evidence for specific null or alternative hypotheses, we performed equivalent non-parametric Bayesian analyses (JASP software (Version 0.8.6.0) (JASP Team, 2018)). The Bayes factor (BF) contrasts the likelihood of the data fitting under the null hypothesis (H0) versus the alternative hypothesis (HA) (Wagenmakers et al., 2018a). For ‘BF10’, values above 1 constitute anecdotal, above 3 moderate, and above 10 strong evidence for the HA (Wagenmakers et al., 2018a, 2018b). To evaluate the evidence for or against an interaction term, we included the main effects in the null model (Wagenmakers et al., 2018b).

## Results

In a TMS (50, 100, 150ms, no-TMS) by stimulus hemifield (left, right) within-subjects design, participants reported, on each trial, on face rotation (clockwise vs counter-clockwise), subjective trustworthiness visibility (1-4 scale), and forced-choice trustworthiness (trustworthy vs untrustworthy) (see Methods). We report the results separately below.

### Rotation task

Figure 2 shows the mean proportions of correct trials over TMS conditions per hemifield. The univariate tests were all non-significant (*p*>0.10). The multivariate tests showed a significant “Hemifield *x* TMS” interaction (*F*(3,14)=3.35, *p*=0.05, η_p_^2^=0.42), in line with our hypothesis.

**Figure 2.**
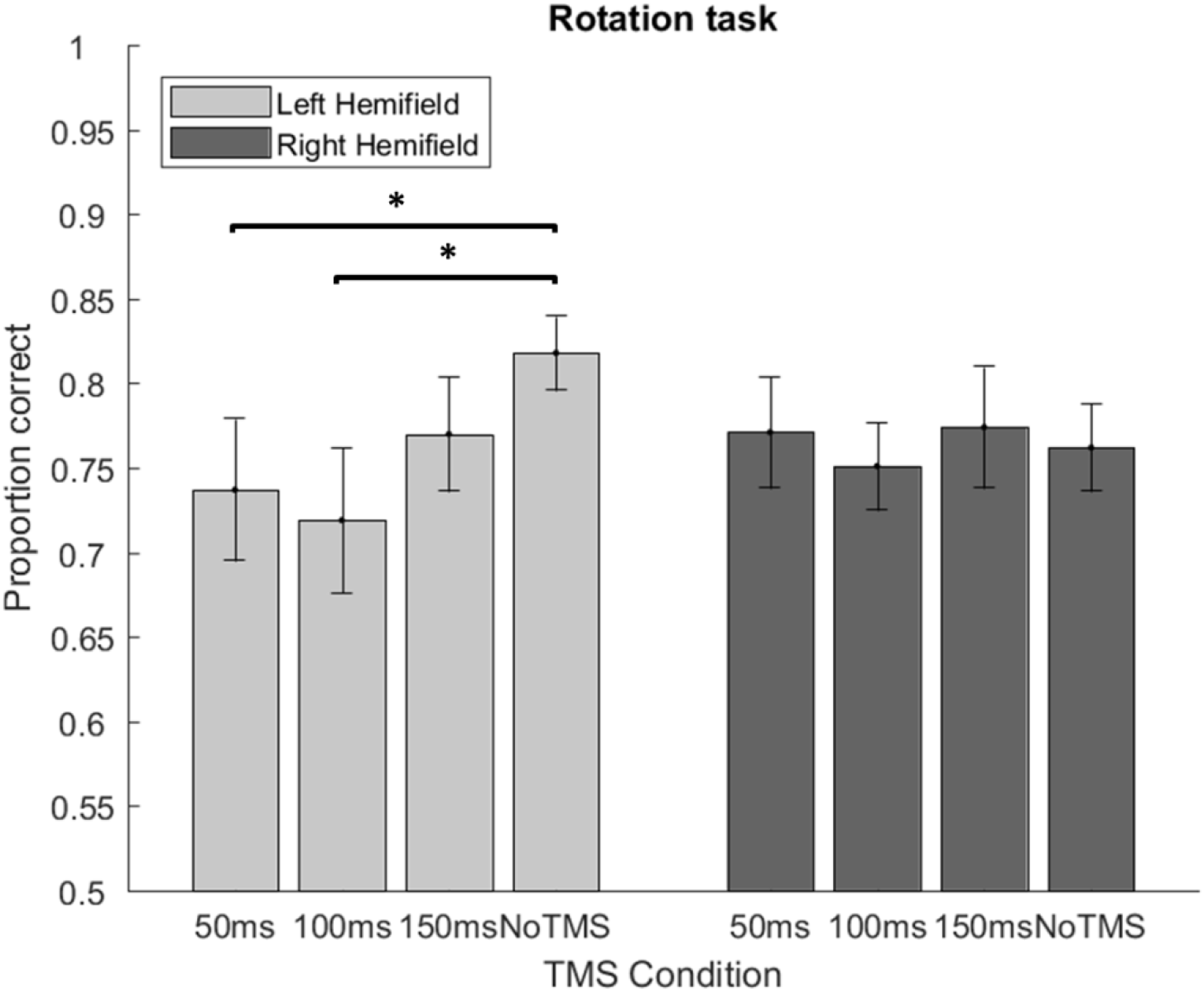
Average accuracies for the rotation discrimination task. Error bars are standard error of the mean (SEM). * corrected *p* < 0.05.

A one-way repeated measures ANOVA univariate follow-up test showed a significant effect of “TMS” on accuracy only for the left hemifield (*F*(2.74, 43.75)=3.85, *p*=0.02, η_p_^2^=0.19). The multivariate test result was non-significant (*F(*3,14)=2.42, *p*=0.11, η_p_^2^=0.34). Rotation discrimination was impaired, as compared to no TMS (*M*=0.82, *SD*=0.09), by TMS at 50 ms (*M*=0.74, *SD*=0.17, *t*(16)=2.53, one-tailed corrected *p*=0.02) and by TMS at 100 ms (*M*=0.72, *SD*=0.18, *t*(16)=4.64, one-tailed corrected *p*=0.02), but not by TMS at 150 ms (*M*=0.77, *SD*=0.82, *p*>0.05).

The above-mentioned results are somewhat inconsistent, since the univariate and multivariate tests support different conclusions. We therefore supplemented our parametric analyses with Bayesian analyses, to investigate how likely the expected effects are given our data. Bayesian analysis provided moderate evidence against a “Hemifield *x* TMS” interaction (BF10=0.22), but provided moderate evidence for a left hemifield TMS effect (BF10=3.41), and supported an impairment of rotation discrimination compared to no TMS at specifically early SOAs (50 ms: BF10=2.91, 100 ms: BF10=3.40, 150 ms: BF10=0.72).

Taken together, these results suggest that rotation discrimination performance is disrupted by TMS to EVC. Though the statistical strength of this result somewhat depends on the tests applied, note that these effects follow precisely the hypothesized pattern based on a large body of previous research: an impairment in rotation discrimination for left hemifield targets at TMS SOAs 50 and 100 ms. This suggests that our TMS protocol could impair forced-choice behavior in our subject sample.

### Subjective visibility task

Figure 3 shows mean subjective visibility ratings per hemifield and TMS condition. The univariate tests showed a significant “Hemifield *x* TMS” interaction (*F*(2.07, 39.35)=3.59, *p*=0.04, η_p_^2^=0.16), but the multivariate tests did not (*F(*3,17)=1.94, *p*=0.16, η_p_^2^=0.26). Both univariate and multivariate follow-up simple effects analyses showed a significant effect of “TMS” only for the left hemifield (respectively *F*(1.46,27.64)=6.16, *p*=0.01, η_p_^2^=0.25; *F*(3,17)=5.11, *p*=0.01, η_p_^2^=0.47). Subjective visibility of trustworthiness was suppressed, as compared to no TMS (*M*=2.46, *SD*=0.10), by TMS at 50 ms (*M*=2.19, *SD*=0.15, *t*(19)=2.53, one-tailed corrected *p*=0.03) and 100 ms (*M*=2.16, *SD*=0.14, *t*(19)=2.94, one-tailed corrected *p*=0.01), but not 150 ms (*M*=2.29, *SD*=0.56, *p*>0.05).

**Figure 3.**
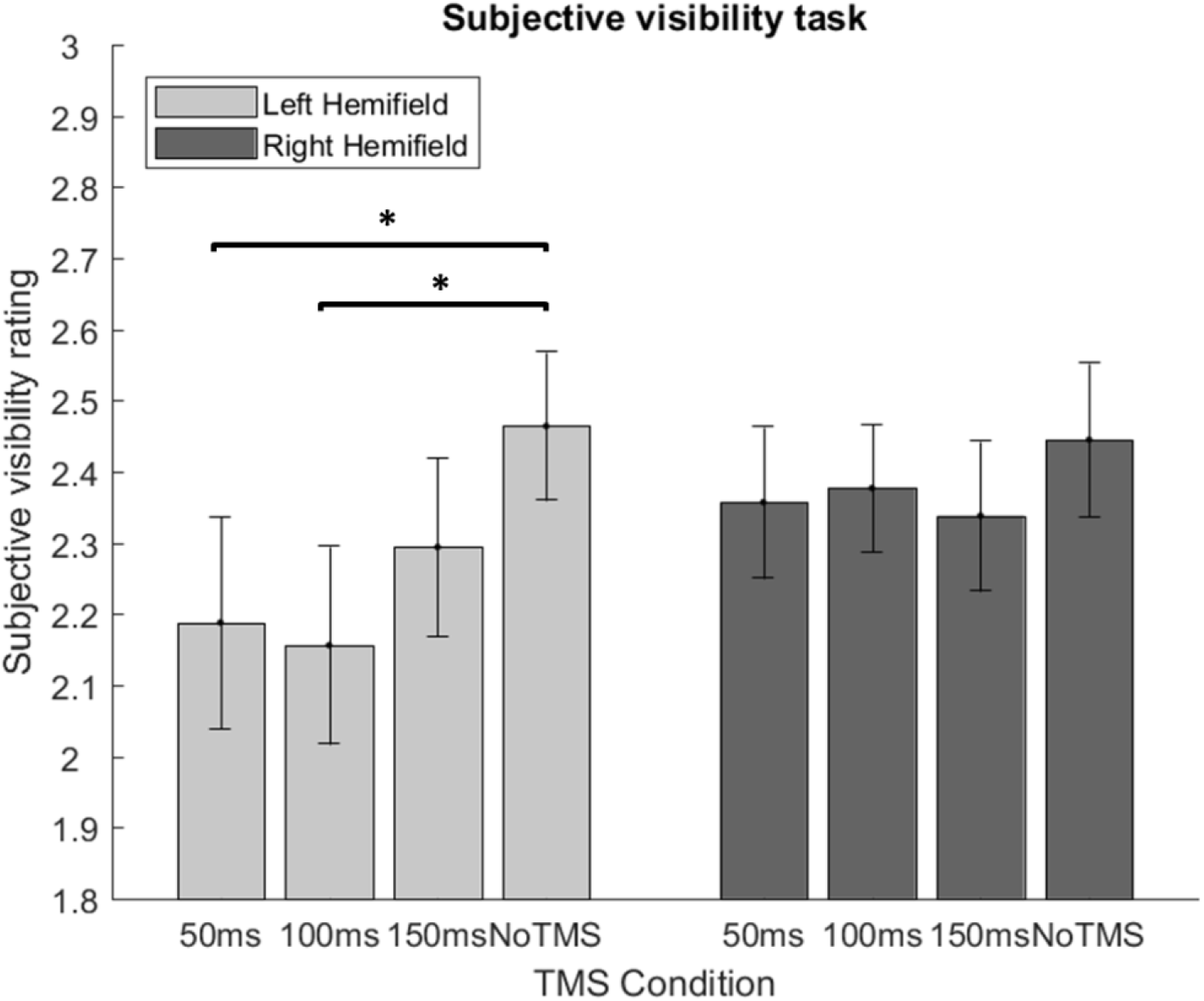
Average subjective trustworthiness visibility ratings on a scale from 1 (“not perceived”) to 4 (“clearly perceived”), with 2 and 3 parametrically in between. Error bars are standard error of the mean (SEM). * corrected *p* < 0.05.

The Bayesian “Hemifield *x* TMS” interaction could not be evaluated due to unacceptable error of the BF estimate. However, Bayesian analysis provided strong support for an effect of TMS specifically for the left hemifield (BF10=29.10). Follow-up analyses provided anecdotal evidence for a suppression effect after TMS at 50 ms (BF10=2.83), moderate evidence for TMS at 100 ms (BF10=6.02), and no evidence for TMS at 150 ms (BF10=0.87). This is in agreement with the previously reported parametric test results. See Supplementary Material for a visualization and details on the proportions of subjective visibility ratings 1, 2, 3 and 4 across TMS conditions and hemifields. Overall, it appears that TMS primarily affected the proportions of trials with lower ratings, e.g. increasing the number of trials in which face trustworthiness was reported as ‘not perceived’ (subjective visibility rating 1).

Altogether, these results show that there was a significant “Hemifield *x* TMS” interaction, due to TMS suppression of specifically left hemifield targets. Subjective visibility ratings were suppressed in the 50 and 100 ms conditions as compared to the no TMS condition. These results suggest that conscious perception of trustworthiness relies on intact EVC at the hypothesized SOAs.

### Trustworthiness task

Figure 4 shows the mean proportions correct on trustworthiness judgments of faces (trustworthy vs untrustworthy) presented in left or right hemifield in different TMS conditions. All univariate and multivariate “Hemifield x TMS interaction” tests were non-significant (*p*>0.10). This suggests there is insufficient evidence to reject any of the null hypotheses. To investigate how much evidence there is *for* a specific null hypothesis, Bayesian statistics (i.e., the Inverse Bayes Factor; BF01) can be used. A two-way Bayesian repeated measures ANOVA showed that the data are 3.49 times more likely under the H0 (i.e., no interaction effect exists) than the HA (i.e., the interaction effect exists). This result gives moderate evidence for the *absence* of a “Hemifield *x* TMS” interaction.

**Figure 4.**
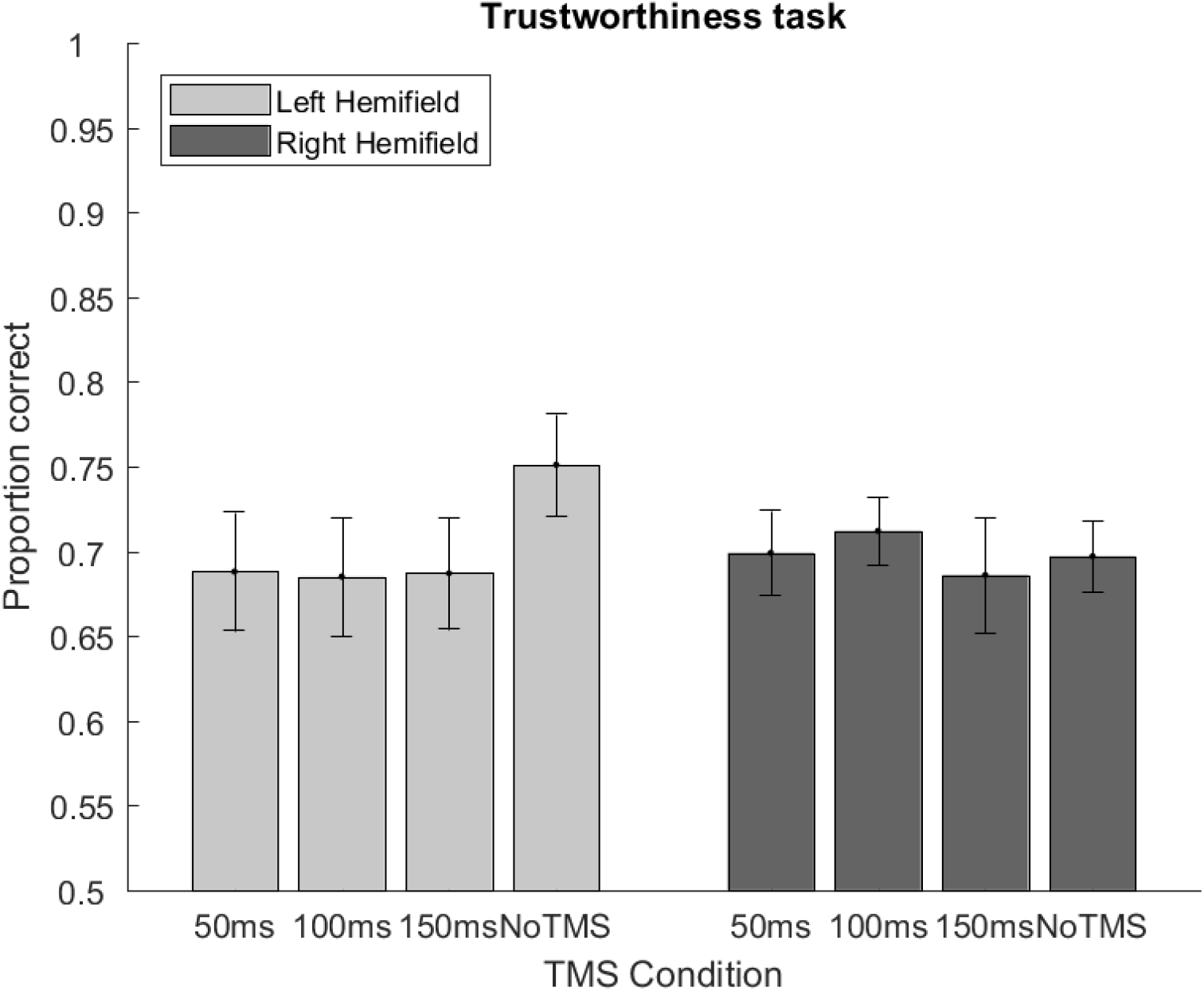
Average trustworthiness discrimination accuracies. Error bars are standard error of the mean (SEM).

In spite of this lack of an interaction, and evidence for absence of such interaction, we evaluated the support for a potential TMS effect on left hemifield targets for full transparency. Univariate and multivariate ANOVAs did not reveal a significant effect of TMS SOA (uncorrected *p*’s: univariate *p*=0.09, multivariate *p*=0.22). According to the Bayesian ANOVA equivalent, our data were equally likely (BF01 = 0.99) under the H0 (i.e., no effect of TMS SOAs) and the HA (i.e., a TMS effect). What support for a TMS SOA exists seems, as can be seen in Figure 4, attributable to the high performance in the no-TMS condition, not to any conventional ‘TMS masking’ pattern over TMS SOAs as can be seen in Figures 2 and 3 (see Discussion below).

Altogether, these results do not convincingly support the notion that trustworthiness discrimination in the left hemifield was impaired by TMS to EVC. This is in agreement with the hypothesis that EVC may not be necessary for forced-choice trustworthiness judgments.

## Discussion

We investigated whether face rotation discrimination, subjective trustworthiness visibility ratings, and face trustworthiness judgments require intact early visual cortex (EVC). We disrupted EVC with double-pulse TMS at different SOAs. At early SOAs (50, 100 ms), TMS to EVC impaired rotation discrimination and suppressed subjective trustworthiness visibility. Yet, there was no evidence for impairment of forced-choice trustworthiness discrimination. These results are in line with the hypothesis that EVC may not be necessary for forced-choice trustworthiness judgments, even though it is required for conscious perception (subjective visibility) of trustworthiness. This suggests that the underlying neural processes can act on our behavior, assessing the trustworthiness of others, without us being aware of it. Although counter-intuitive, such automatic processes have also been shown for other types of information (see below). While conscious processing allows reflection and flexibility, for ecologically relevant information quick and automatic processes may be at least as important. Interestingly, the intrinsically social process of trustworthiness evaluation might fall in this category, further highlighting its ecological relevance.

Our conclusions converge with several previous reports. For instance, in 7-month-old infants, differences in event-related potentials (ERPs) elicited by unconscious (un)trustworthy faces were found for central and frontal but not occipital electrodes (Jessen & Grossmann, 2017). Moreover, the amygdala and insula but not the EVC were differentially activated by (un)trustworthy faces during an implicit trustworthiness discrimination task (Winston, Strange, O’Doherty, & Dolan, 2002). Furthermore, unconscious trustworthiness evaluation affected amygdala activity but did not affect activity in fusiform areas (Freeman et al., 2014). This resembles the mechanisms assumed to underlie unconscious emotion processing, where several studies suggest that strong emotional signals such as facial expressions can influence amygdala activation via a subcortical route in the absence of conscious awareness (Celeghin, de Gelder, & Tamietto, 2015; Tamietto & de Gelder, 2010). In emotional expressions, single features such as wide eyes are sufficient to elicit emotion-related differences in neural activation (Whalen et al., 2004). Crucially, though, face trustworthiness is characterized by more complex feature arrangement (Todorov et al., 2015). Hence, the question arises to what degree subcortical mechanisms are sufficient to evaluate the complex feature arrangements necessary to differentiate face trustworthiness.

The debate about whether the neural pathway underlying unconscious face trustworthiness processing is entirely sub-cortical, is ongoing. In prior work, two higher order visual areas, namely the superior temporal sulcus (STS) and the fusiform gyrus, were differentially activated by unconscious (un)trustworthy faces (Winston et al., 2002). However, the STS contrast was mostly seen during an explicit task, and the fusiform gyrus contrast might be explained by modulatory feedback responses from the amygdala (Morris et al., 1998). There is some indirect evidence *against* a solely sub-cortical route. While sub-cortical emotion processing is assumed to involve predominantly low spatial frequencies, i.e. coarse information (Vuilleumier, Armony, Driver, & Dolan, 2003), amygdala activation was modulated by face trustworthiness for both images containing only low and images containing only high spatial frequencies (Said, Baron, & Todorov, 2009). Future studies are necessary to elucidate this issue.

Faces seem to be processed differently between the two cerebral hemispheres. Previous studies have established the right hemispheric dominance for face perception in general (De Heering & Rossion, 2015; Gainotti & Marra, 2011; Pitcher, Walsh, Yovel, & Duchaine, 2007; Rangarajan et al., 2014; Rossion, Hanseeuw, & Dricot, 2012; Van Belle et al., 2011). More specifically, some evidence suggests that especially the right hemisphere is involved in the processing of emotional expressions (Demaree, Everhart, Youngstrom, & Harrison, 2005) and face trustworthiness (Dzhelyova, Perrett, & Jentzsch, 2012; Okubo, Ishikawa, & Kobayashi, 2013). In line with this strong right-lateralization of face processing, we targeted only the right EVC. However, the right hemisphere is not exclusively involved in this type of processing. For instance, there is some evidence suggesting that the left vs right hemispheres are preferentially processing positive and negative emotions, respectively (Okubo, Ishikawa, & Kobayashi, 2017). Moreover, a recent study found that faces presented in the right visual field (and thus processed by the left hemisphere) were rated as more trustworthy compared to faces presented in the left visual field (Slepian, Young, & Harmon-Jones, 2017). It thus remains unclear what is the exact role of the left EVC in the processing of face trustworthiness.

Although our analyses did not support a TMS effect on trustworthiness processing, it is apparent from Figure 4 that performance in no-TMS trials in the left hemifield was higher than in the three TMS conditions. We have previously discussed the possible complications of no-TMS trials in randomized event-related experimental TMS designs; the unexpected omission of pulses on trials can be surprising, with varying consequences on performance (De Graaf & Sack, 2010; De Graaf et al., 2011; Duecker & Sack, 2013). This, the lack of statistical support for TMS effects, and the fact that there was no difference in performance across the three TMS SOAs at all, leads us to the conclusion that trustworthiness judgments were unimpaired by TMS in our experiment. The alternative interpretation is that there was a general decrease in trustworthiness discrimination performance after TMS, as compared to no-TMS. There is some evidence suggesting that EVC may be functionally relevant for face processing for a longer period of time. We previously observed delayed or prolonged TMS suppression of face stimuli (De Graaf et al., 2012), though not with SOAs as late as 150-200 ms. There are reports of TMS masking at later (>150ms) SOAs for more complex tasks such as categorization of natural scenes (Camprodon, Zohary, Brodbeck, & Pascu, 2009; Koivisto, Railo, Revonsuo, Vanni, & Salminen-Vaparanta, 2011), figure-ground segregation (Heinen, Jolij, & Lamme, 2005), and complex visual search (Dugué, Marque, & VanRullen, 2011; Juan & Walsh, 2003). Still, a temporally unspecific TMS masking effect seems unlikely.

A general decrease in performance after TMS could be due to the fact that we used connecting SOAs (see Methods), limiting the effective temporal resolution (De Graaf et al., 2014). Even though we used double-pulse TMS because of its stronger impact on EVC as compared to single TMS pulses, this difference is small (Gerwig, Niehaus, Kastrup, Stude, & Diener, 2005; Kammer & Baumann, 2010; Ray, Meador, Epstein, Loring, & Day, 1998). Future studies might replicate the current findings with higher temporal specificity and a broader range of SOAs. Another limitation of the current report involves the unusually large size of our visual stimuli. Usually in occipital TMS suppression studies, stimuli are small so as to fully suppress them with TMS pulses. However, our goal here was not to make the stimuli invisible, but to disrupt specific aspects of their processing, including rotation judgments and the subjective visibility of trustworthiness (as opposed to visibility of the face itself). The magnitudes of effect here do not deviate strongly from those reported in previous work. Lastly, we designed and analyzed this project with the explicit a priori determined approach of evaluating separately potential effects on rotation judgment, 2AFC trustworthiness judgment, and subjective visibility of trustworthiness. Future studies could alternatively evaluate blindsight-like performance for trustworthiness information more directly, by asking per trial whether trustworthiness was perceived and evaluating 2AFC judgments on specifically unseen trials (Boyer et al., 2005; Jacobs et al., 2012; Jolij & Lamme, 2005; Koenig & Ro, 2018; Ro, Shelton, Lee, & Chang, 2004; Christensen et al., 2008).

Finding no TMS suppression of trustworthiness judgment is a null result. Generally, null results in TMS are difficult to interpret (De Graaf & Sack, 2010), since the complex combination of TMS stimulation parameters and stimulus characteristics determines whether effects are detected or not. Perhaps different or individualized cortical targeting, stimulation intensity, or face stimulus duration procedures would impair trustworthiness judgment after all. We recently presented a taxonomy for null result interpretation, with guidelines on design and interpretation (De Graaf & Sack, 2018). In the current study, we included a ‘neural efficacy check’ (rotation task effects) and Bayesian support for the null finding (Wagenmakers et al., 2018a, 2018b). This makes the collection of results on the rotation task, subjective visibility, and forced-choice trustworthiness judgments, of interest. The positive support for a role of EVC in rotation and conscious trustworthiness processing, and negative result for forced-choice trustworthiness processing, were all in accordance with the hypothesis and prior research.

## Conclusion

TMS is ideally suitable to investigate whether and when specific cortical areas are necessary for certain processes (Pascual-Leone, Walsh, & Rothwell, 2000). We extend the existing literature by showing that conscious perception of face trustworthiness can be, but forced-choice trustworthiness judgments may not be, disrupted by TMS to EVC. Our results regarding the EVC, the lowest-level cortical area in the visual hierarchy, pave the way for a more in-depth exploration of the underlying neural pathways. A fruitful next step would be to investigate the relevance of higher order cortical visual areas such as the STS for the processing of face trustworthiness. Moreover, it would be highly interesting to study which subcortical areas are involved. Although it is not possible to directly stimulate subcortical regions with TMS, they could be indirectly stimulated via cortical areas (Wagner, Rushmore, Eden, & Valero-Cabre, 2009; Wang et al., 2014). Lastly, TMS could be used in combination with neuroimaging methods such as fMRI and EEG, to visualize the entire network underlying the processing of face trustworthiness.

## Supporting information

Supplementary Analysis

## Acknowledgements

This work was supported by the Netherlands Organization for Scientific Research (NWO: VENI to TG [grant number 451-13-024], VICI to AS [grant number 453-15-008]).

