## Supplementary Analysis for "Can processing of face trustworthiness bypass early visual cortex? A transcranial magnetic stimulation masking study"

### Supplementary material: proportion of subjective visibility ratings across conditions

We found decreased subjective trustworthiness visibility ratings in the left hemifield after TMS at 50 ms and 100 ms (see ‘Subjective visibility task’ under ‘Results’). The question arises where this suppressive effect of TMS is coming from: does TMS only influence trustworthiness visibility in the higher part of the scale, having an effect on the clarity of perception? Or does TMS also have an effect in the lower part of the scale, increasing the number of trials in which trustworthiness is not perceived at all? To investigate how TMS affected subjective trustworthiness visibility, we calculated the proportion of trials in each level of the 4-point scale as a function of TMS and hemifield. Inspection of Supplementary Figure 1 shows that the proportion of trials in which participants indicated not having perceived the trustworthiness at all (visibility rating 1) seems to be increased specifically for the left hemifield 50 and 100 ms conditions. However, no significant differences in proportions were found across conditions when comparing them in a 4 (visibility rating) x 4 (TMS) x 2 (hemifield) repeated-measures ANOVA.

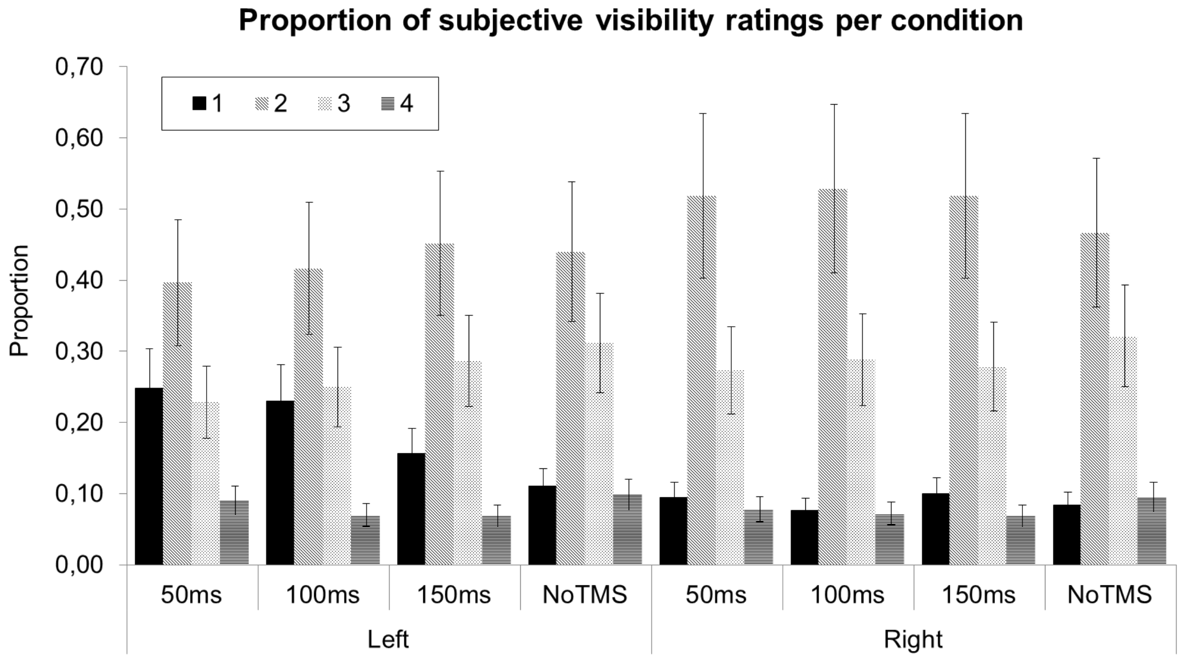

*Supplementary Figure 1*. Proportion of trials in each level of the 4-point subjective trustworthiness visibility scale, separately per TMS condition and hemifield. For each condition, the leftmost bar indicates the proportion of trials with subjective visibility rating 1 (trustworthiness "not perceived") and the rightmost bar indicates the proportion of trials with subjective visibility rating 4 (trustworthiness "clearly perceived"), with rating 2 and 3 in between. Error bars are standard error of the mean (SEM).
